# Novel midgut smooth muscle necrosis (MSMN) in translucent or glass post-larvae of whiteleg shrimp

**DOI:** 10.1101/2025.07.05.663262

**Authors:** Jiraporn Srisala, Piyachat Sanguanrut, Sorawit Powtongsook, Krit Khemayan, Kallaya Sritunyalucksana, T.W. Flegel

## Abstract

From the Asia-Pacific region, the suspected TPD/GPD specimens of postlarvae (PL) preserved in Davidson’s fixative and 95% ethanol were sent to our laboratory to prepare for PCR and histological analysis. Interestingly, PCR tests with the specimens were negative with the V_*P*HLVD_(*vtcc*2 and *vtcc*3 gene) primers, but positive with the V_*P*TPD_(*vhvp*-2 gene) primers. There were no characteristic lesions of acute hepatopancreatic necrosis disease detected in these specimens. However, in the sections of several specimens from two affected tanks, the midgut was cut tangentially in the abdominal region, revealing a novel histopathology characterized by severe necrosis specifically in the thin, smooth muscle tissues underlying the midgut lumen epithelium. This comprised nuclear pyknosis and karyorrhexis limited to the smooth muscle and occasionally affecting the midgut epithelium but not involving the surrounding skeletal muscles. The pathology is called midgut smooth muscle necrosis (MSMN). Such pathology would undoubtedly lead to loss of midgut peristalsis and cessation of feeding, but it can be easily overlooked without careful histological examination using a high magnification microscope objective. No bacterial cells were evident, suggesting that the lesions arose from a virus or from a specific toxin response. This report is an urgent plea for colleagues studying TPD/GPD to review their samples histologically to help in determining whether MSMN is pathognomonic for TPD/GPD.

## INTRODUCTION

Newly emerging diseases causing high mortality of post-larvae of *Penaeus vannamei* in China since 2019 include translucent post-larva disease (TPD) and glass post-larval disease (GPD) that have been reported by Zou *et al.* (2020), Yang *et al.* (2022) and Xu *et al.* (2023). The etiologies of these diseases have been reported to be either toxin-producing strains of *Vibrio parahaemolyticus* (VP) or a virus in the family *Marnaviridae* named *Baishivirus*. Two virulent strains of *Vibrio* were isolated. The one causing TPD was named *V. parahaemolyticus* TPD (*Vp*_TPD_) by Zou *et al.* (2020), while the one causing GPD was named highly lethal *Vibrio* disease (HLVD) Vp (*Vp*_HLVD_) by Yang *et al.*, 2022). The toxin from *Vp*_TPD_ (Liu *et al.*, 2023) was called Tc toxin while that produced by *Vp*_HLVD_ (Yang *et al.*, 2023) was called *vhvp*-2 toxin. The clinical symptoms caused by VP_AHPND_ and *Vp*HLVD are similar (Zou *et al.*, 2020; Yang *et al.*, 2022; Tran *et al.*, 2013) with the infected shrimp demonstrating hepatopancreatic necrosis with sloughing of the epithelial cells of hepatopancreatic tubules and midgut (Sirikarin *et al.*, 2015), similar to the pathology of acute hepatopancreatic necrosis disease (AHPND)(Tran et al. 2013) caused by strains of *V. parahaemolyticus* that produce PirA and PirB toxins (Sirikharin et al., 2015) that differ from Tc toxin and *vhvp*-2 toxin. Another publication by Xu *et al.* (2023) described glass post larvae disease (GPD) and proposed that the causal agent was a new RNA virus called *Baishivirus* (Family *Marnaviridae*). However, the histopathology associated with GPD, including that in the moribund shrimp challenged with inoculum containing *Baishivirus* has not been described.

From a country in the Asia-Pacific region in early 2024, we received specimens of the whiteleg shrimp *P. vannamei* described as exhibiting gross signs of “translucent post-larvae disease” (TPD) (Liu *et al.*, 2023; Zou *et al.*, 2020) or “glass post-larvae disease” GPD (Xu *et al.*, 2023). The specimens were analyzed for possible causes of their translucence by using PCR methods to detect *vhvp*-2 toxin and Tc toxins (*vtcc2* and *vtcc3*), as well as histological examination to determine the possible presence of hepatopancreatic necrosis described as the pathology for TPD or perhaps a specific histopathology that might be associated with GPD.

## METHODS

### Samples

The PL9 samples originated from 2 hatchery tanks from a single facility that exhibited PL translucency and mortality. The PL were collected in 95% ethanol for DNA/RNA extraction or in Davidson’s fixative for histological examination with H&E staining by standard methods (Bell and Lightner, 1988). The number of 10 PL preserved in 95% ethanol were pooled into 1 sample from each hatchery tank for DNA and RNA extraction. The histological samples were altogether 38 specimens, 20 from hatchery Tank A and 18 from hatchery tank B, and they resulted in 2 paraffin blocks for each tank (n= 9-10 each slide).

### DNA and RNA extraction

The eyestalks of PL were removed prior to the DNA extraction (PCR for TPD and AHPND) and RNA extraction (RT-PCR for *Baishivirus*) and homogenized in DNA lysis buffer (50 mM Tris-base (pH 9.0), 100 mM EDTA, 50 mM NaCl, 2% (w/v) SDS and 100 μg/ml proteinase K) or Trizol reagent (Invitrogen, USA). DNA extraction employed Exgene™ cell SV mini kits (GeneAll, Korea) according to the manufacturer’s instructions. RNA extraction followed the Trizol reagent protocol. The RNA pellet was re-suspended with DNase/RNase free water and digested with DNase I (NEB, USA) following the manufacturer’s protocol. The DNA extracts and RNA extracts were subjected to PCR and RT-PCR detection, respectively.

### PCR detection of TPD and AHPND

For the PCR method, four primer sets were used. These were TcC2*-*F/R for *vtcc2* gene and TcC3*-*F/R for *vtcc3* gene (Yang *et al.* 2023) and *vhvp-2*-F1/R1 and *vhvp-2*-F2/R2 (Liu *et al.* 2023). The *vhvp-2*-F1/R1, and *vhvp-2*-F2/R2 methods targeted different regions of *vhvp-2* gene of *Vp*_TPD_. The primer sequences are shown in Table 1 with the detailed PCR protocols and the expected amplicon sizes. PCR reactions were performed in a 12.5 μl mixture containing 1× OneTaq Hot Start Master Mix (NEB, USA), 0.4 μM each primer and 50 ng of DNA template. Also included was PCR detection for VP_AHPND_ using the AP4 method (Table 1) (Dangtip *et al.*, 2015).

**Table 1.**
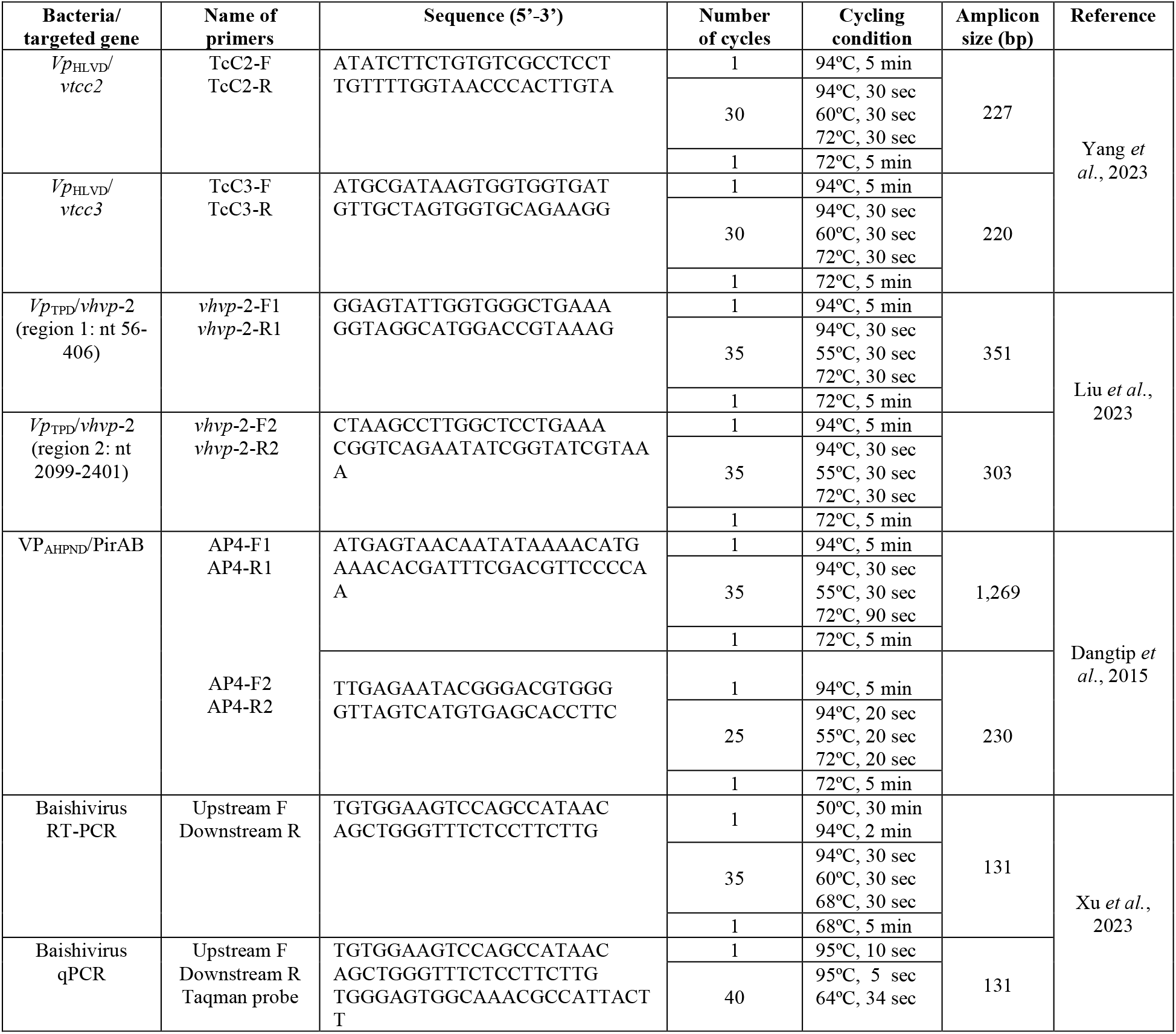
Information for the PCR/RT-PCR methods for TPD/GPD and AHPND detection

### RT-PCR and qPCR Detection of *Baishivirus*

For *Baishivirus*, the RT-PCR method of Xu *et al.*, 2023 was used and the primers are listed in Table 1. The RT-PCR reaction was performed in a 12.5 μl mixture containing 1 Reaction Mix (Invitrogen, USA), 0.4 μM each of upstream primer F and downstream primer R, 0.5 μl of SuperScript III RT/Platinum Taq Mix (Invitrogen, USA) and 50 ng of RNA template.

For qPCR analysis, 50 ng of cDNA template was added to a master mix that consisted of 1× qPCRBIO Probe Mix Lo-Rox (PCR Biosystems, UK), 0.4 μМ of upstream F and downstream R primers, 0.2 μМ TaqMan probe (IDT, Singapore) as shown in Table 1. The thermal cycling protocol is also shown in Table 1.

### *In situ* hybridization (ISH) testing

In this study, a *Baishivirus* fragment template targeting the ORF2 gene (GenBank: ON550424.1) was synthesized by GenScript (GenScript, USA). To prepare the probe, new primers were designed to extend the target region and include the amplicon previously reported by Xu *et al.* (2023). Briefly, the primers 573F: TTACGTTGCT GGGAAGGTCA) and 573R: TGGAGAGGAAGTTCACGGAC were used to generate a DIG-labeled DNA probe (amplicon size: 573 bp) for *Baishivirus* detection using a PCR DIG-labeling kit (Roche, Germany). The probe was then purified using a Gel/PCR Cleanup Kit (Geneaid, Taiwan), following the protocols provided by the manufacturer.

The protocol for ISH has been previously described (Sanguanrut *et al.*, 2022). Briefly, slides containing adjacent tissue sections were de-paraffinized, rehydrated in TNE buffer (500 mM Tris-Cl, 100 mM NaCl, l0 mM EDTA) before treatment with 5 µg proteinase K (Sigma, USA). The prepared tissue sections were incubated with 400 ng of a DIG-labeled *Baishivirus* probe (per slide) in hybridization buffer at 42 ºC overnight. After incubation the slides were washed and incubated with blocking buffer and further incubated with 1:500 anti-DIG-AP antibody under dark conditions. Finally, the hybridized tissue sections were washed and developed for signal using NBT/BCIP solution (Roche, Germany), followed by counterstaining with 0.5% Bismarck Brown Y (Sigma, USA) for microscopic examination (Leica DM750, ICC50W model with a digital camera and LASV4.12 Software). Unfortunately, we did not have any positive control shrimp samples.

## RESULTS

### PCR testing gave negative test results for AHPND and TPD

Samples from the 2 hatchery tanks gave negative PCR test result for VP_AHPND_ toxin A by the AP4 method (Data not shown). For PCR detection of TPD toxin genes, specimens from both tanks gave positive results with 2 targets of the *vhvp*-2 toxin gene (regions 1 and 2) (**Fig. 1A and 1B**), but not with the other two targets *vtcc2* or *vtcc3* (regions 2 and 3) **Fig. 1 B and C**).

**Figure 1.**
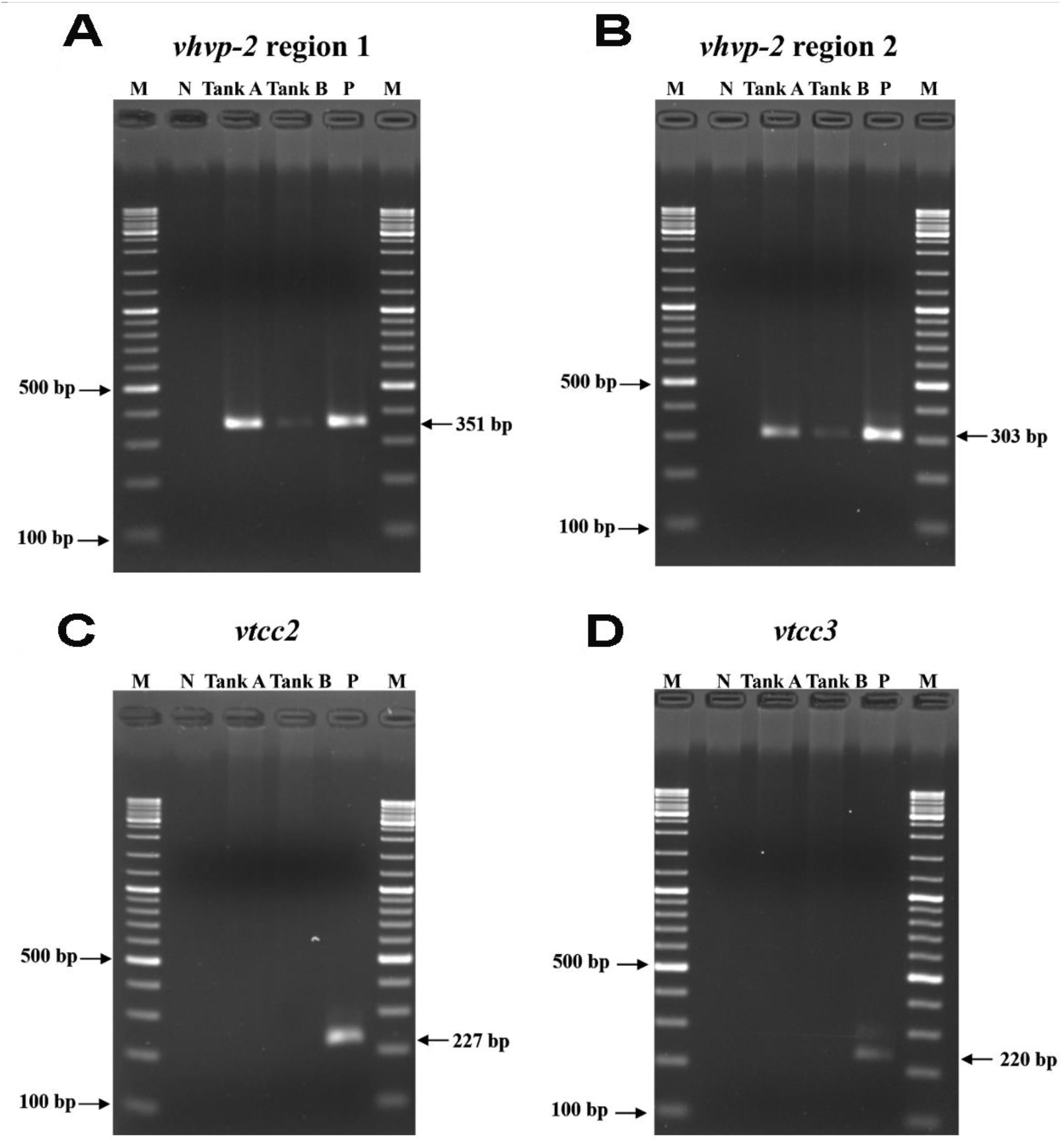
**A&B.** Positive agarose gel electrophoresis results for *vhvp*-2 gene regions 1 and 2. **C&D.** Negative gel electrophoresis results for *vtcc*2 gene regions 3 and 4.

### RT-PCR and qPCR testing for *Baishivirus* gave negative test results

Our RT-PCR and qPCR testing using the method recommended for detection of *Baishivirus* by Xu *et al.* (2023), gave negative findings by RT-PCR and gel electrophoresis (**Fig. 2**) consistent with qPCR analysis (data not shown) and validated through ISH assays. However, it is possible that the samples we received had been in transit too long or had been improperly prepared for RNA preservation, leading to negative test results for *Baishivirus* detection.

**Figure 2.**
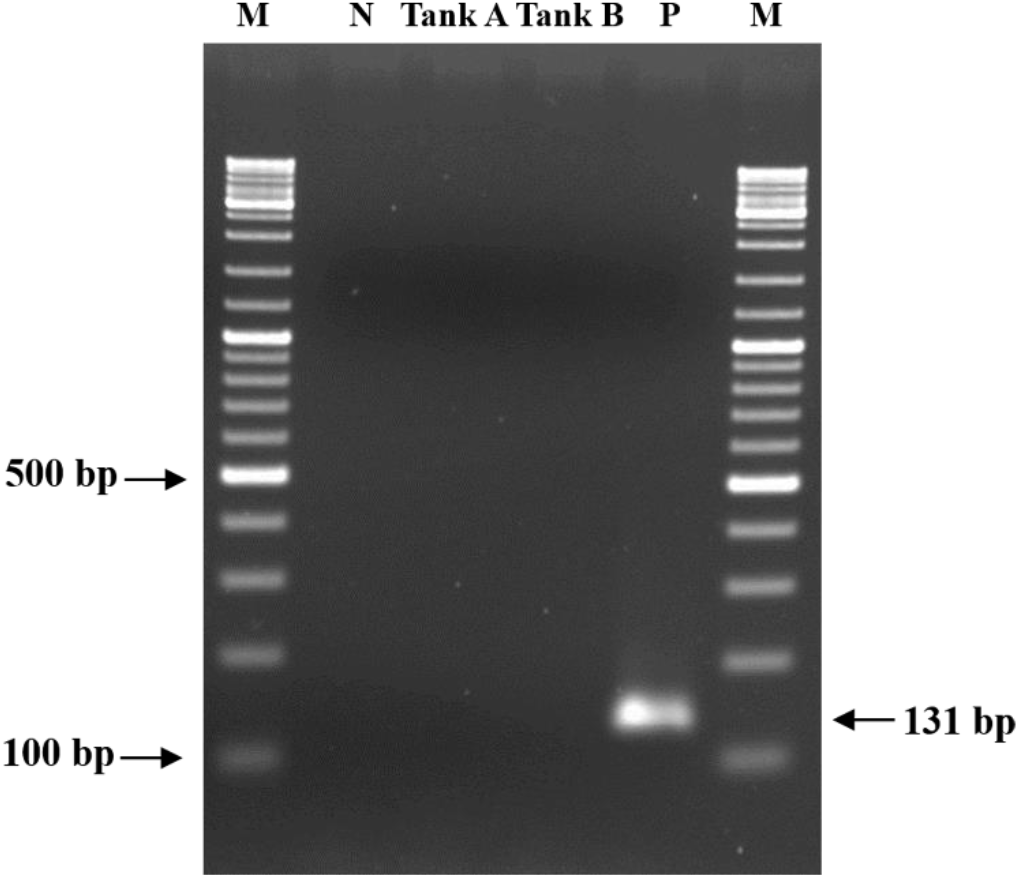
Agarose gel electrophoresis analysis of Baishivirus. M = 2 log DNA marker; N = Negative control; P = Positive control; Tank A= RNA specimen from hatchery tank A and Tank B: RNA specimen from hatchery tank B.

### No evidence for pathology in the hepatopancreas

Based on a previous publication (Zou *et al.*, 2020) regarding TPD, our initial examination of the PL specimens focused on their hepatopancreas (HP). The aim was to find evidence for extensive sloughing of tubule epithelial cells with or without presence of bacteria, like the pathology caused by AHPND (Tran *et al.*, 2013). Of the total of 38 PL tissue sections, hepatopancreatic tissue was included in the tissue sections of 13 of the 20 specimens from hatchery Tank A and 11 of 18 from hatchery Tank B. The total of 24 PL sections showed normal HP histology. Four representative examples are illustrated in **Fig. 3** for specimens from Tank A. These 4 specimens were selected because they also showed novel histopathology in midgut smooth muscle that is subsequently described herein. In Fig. 3, the representative hepatopancreatic tissues show no signs of tubule epithelium sloughing, hemocytic aggregation or bacterial cells that are characteristic of AHPND (Tran *et al.*, 2013). Thus, our observations did not reveal the HP pathology of TPD described by Zou *et al.*, 2020.

**Figure 3.**
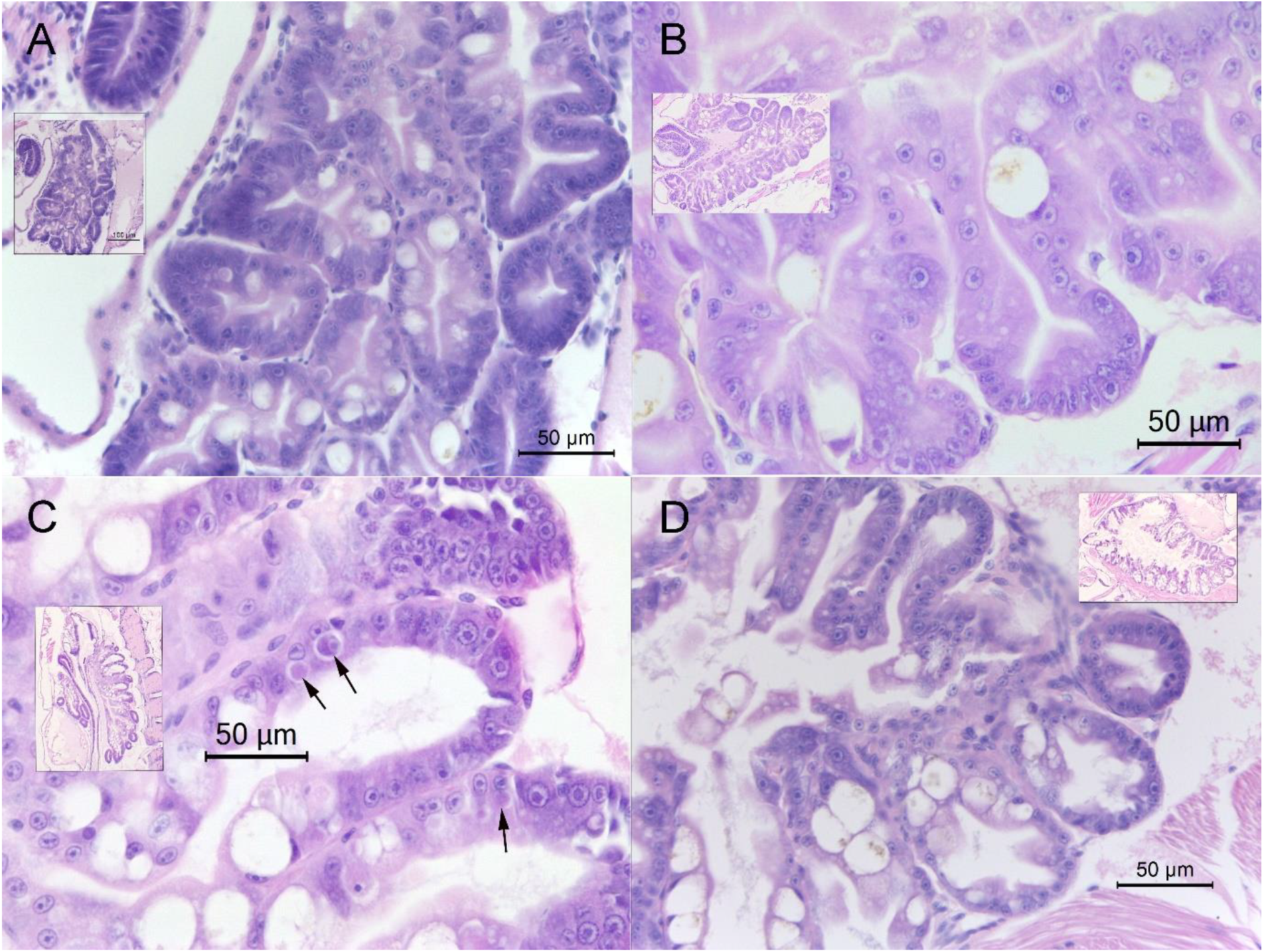
Histology of hepatopancreatic tissue of 4 specimens from hatchery Tank A. These are representative samples for the HP of all 24 PL specimens from hatchery Tanks A and B whose sections showed HP tissue. They all showed normal histology without HP tubule cell sloughing, hemocytic aggregation or bacterial colonies associated with AHPND pathology. These are magnified portions of the whole HP tissue section that is shown in a small inset and they correspond to the HP seen in whole shrimp sections in Figs. 5,6, 7 and 8 respectively. Note that some of the cells (arrows) in the HP in (C) show lesions typical of *Penaeus vannamei* solinvivirus (PvSV) (arrows).

Although the 4 representative specimens from Tank A showing normal HP histology in Fig. 3, they showed novel abdominal midgut pathology that has not been previously described from shrimp and will be described in detail below. Of the 13 specimens with HP from Tank A, tissue sections of 12 included abdominal midgut tissue and 7 exhibited this novel midgut pathology while the remainder were normal. The 11 sections with HP from Tank B, only 8 included abdominal midgut tissue and only 1exhibited this novel pathology. No other histopathology was observed in the total of 38 samples examined from the 2 hatchery A and B tanks.

### Novel histopathology confined specifically to midgut smooth muscles

The same 4 specimens used to prepare Fig. 3 also showed very unusual histopathology only in the smooth muscle tissue of the mid-abdominal region the midgut. To understand this unusual histopathology, the photomicrographs of normal hepatopancreatic and abdominal midgut histology, particularly focused on the smooth muscular layer of the midgut are shown in **Fig. 4**. This layer underlies the midgut lumen epithelium in 2 layers of smooth muscle cells, an inner layer with muscle fibers that encircle the midgut epithelium and an adjacent, outer layer with fibers that extend longitudinally along the midgut. Together, these muscles interact to achieve peristalsis.

**Figure 4.**
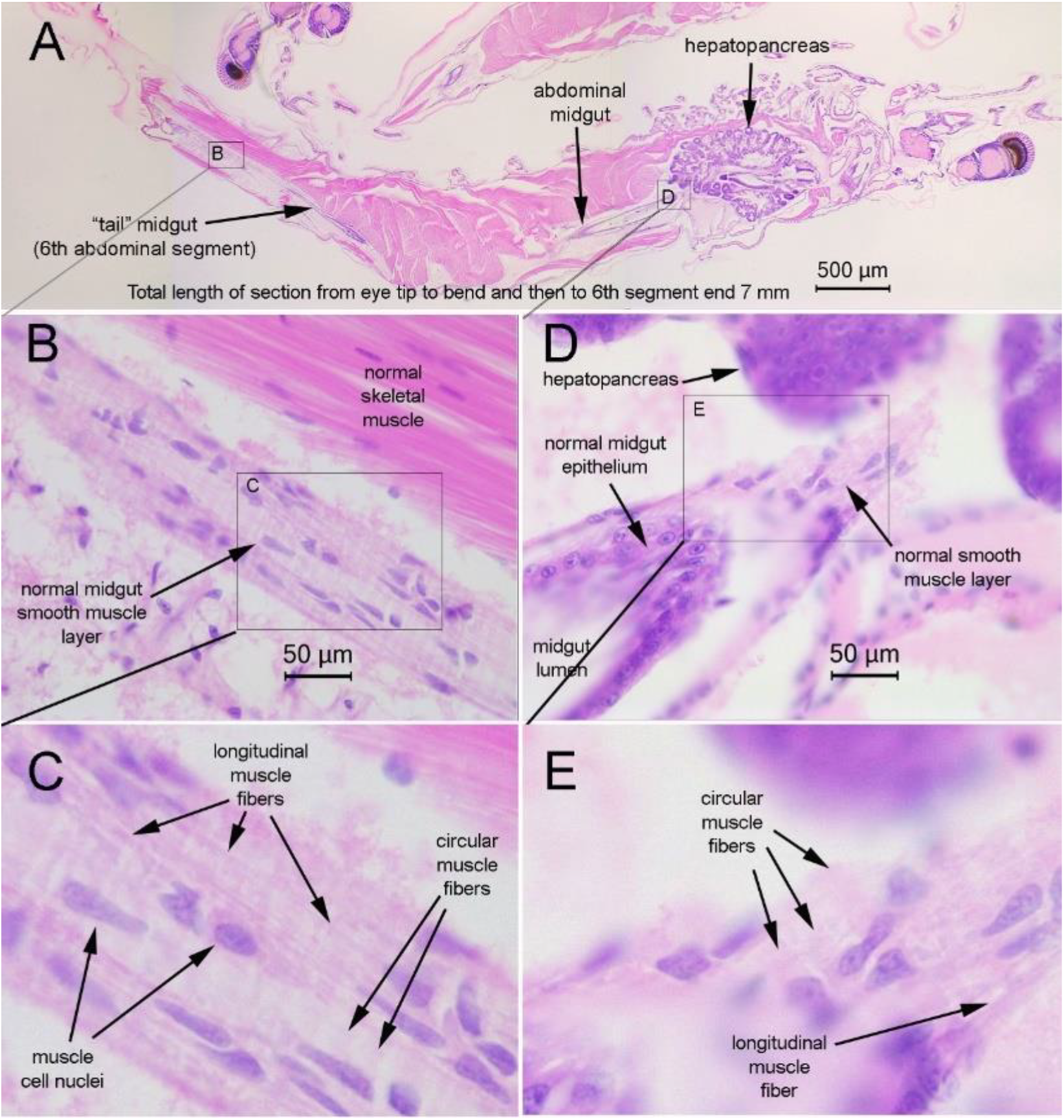
Example of normal midgut histology from a PL specimen from hatchery Tank A. This tissue section includes the extended 6^th^ abdominal segment (“tail”). (A) Low magnification of the whole PL. (B) medium and (C) high magnification of a lateral cut of the midgut in the 6^th^ abdominal segment and showing normal smooth muscle cell nuclei plus normal circular and longitudinal fibers. (D) medium and (E) high magnification of the midgut showing similar normal histology in the midgut region near the hepatopancreas.

To see the novel pathology in the midgut smooth muscle layers clearly by microscopic examination, a high magnification objective of 40 to 100 times is needed. Because the midgut layers are very thin, the best view is obtained only when the midgut is cut tangentially to provide a slanted view that shows a portion of the midgut lumen, a portion of the epithelium that surrounds the lumen, and then 2 smooth muscle layers that are sandwiched between the midgut epithelium and the midgut sheath.

In comparison to Fig. 4, **Fig. 5** shows sections of PL specimens from the same hatchery Tank A as the normal specimen shown in Fig. 3. It includes a tangential section such as described above that passed through the midgut within the region of the first 5 abdominal segments.

**Figure 5.**
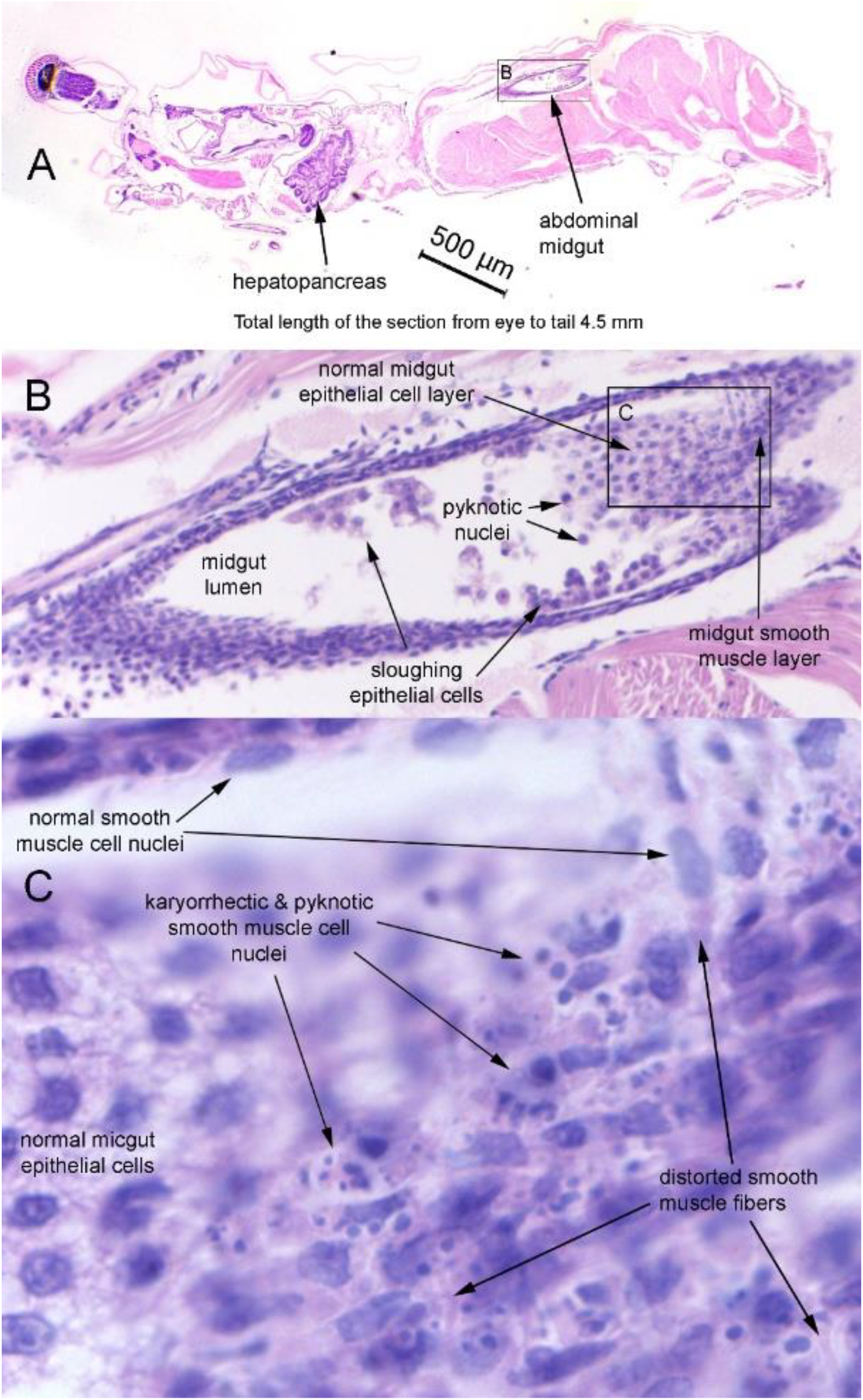
PL from Tank A showing novel necrosis of the smooth muscle layer in the midgut in the region of the first 5 abdominal segments. The 6^th^ abdominal section is not within the cut. (**A**) Low magnification with a box indicating the enlarged region in B. (B) Shows areas of normal midgut epithelial cells together with those sloughed and with pyknotic nuclei. The box indicates the enlarged region shown in C. (C) Clearly shows massive necrosis of the smooth muscle layer including pyknotic and karyorrhectic nuclei together with distorted smooth muscle fibers. This contrasts sharply with the normal histology shown in Fig. 4B to E and is not easily visible at the magnifications in 5A and B.

The 6^th^ abdominal (“tail’) segment was not included in the cut, and it would have added approximately 30% (1.4 mm) to the total length of the specimen. **Fig. 5C** clearly shows massive necrosis of the smooth muscle layers that exhibit extensive, basophilic, nuclear pyknosis and karyorrhexis together with distorted, eosinophilic smooth muscle fibers that resemble cooked spaghetti noodles. We refer to this pathology as midgut smooth muscle necrosis (MSMN). A high magnification of a portion of the normal HP of this same specimen is shown in Fig. 3.

In addition, **Figure 6** shows MSMN histopathology from a second specimen from hatchery Tank A. The section cut is lateral to the abdominal region of the midgut and demonstrates the difficulty in easy diagnosis of the MSMN in such cuts due to the thinness of the muscle layer that underlies the midgut. Distortion of the muscle fibers cannot be clearly seen, and the severity of the necrosis is not as easily detected as in the tangential section seen in Fig. 5C. Note that many midgut epithelial cells have sloughed into the midgut lumen.

**Figure 6.**
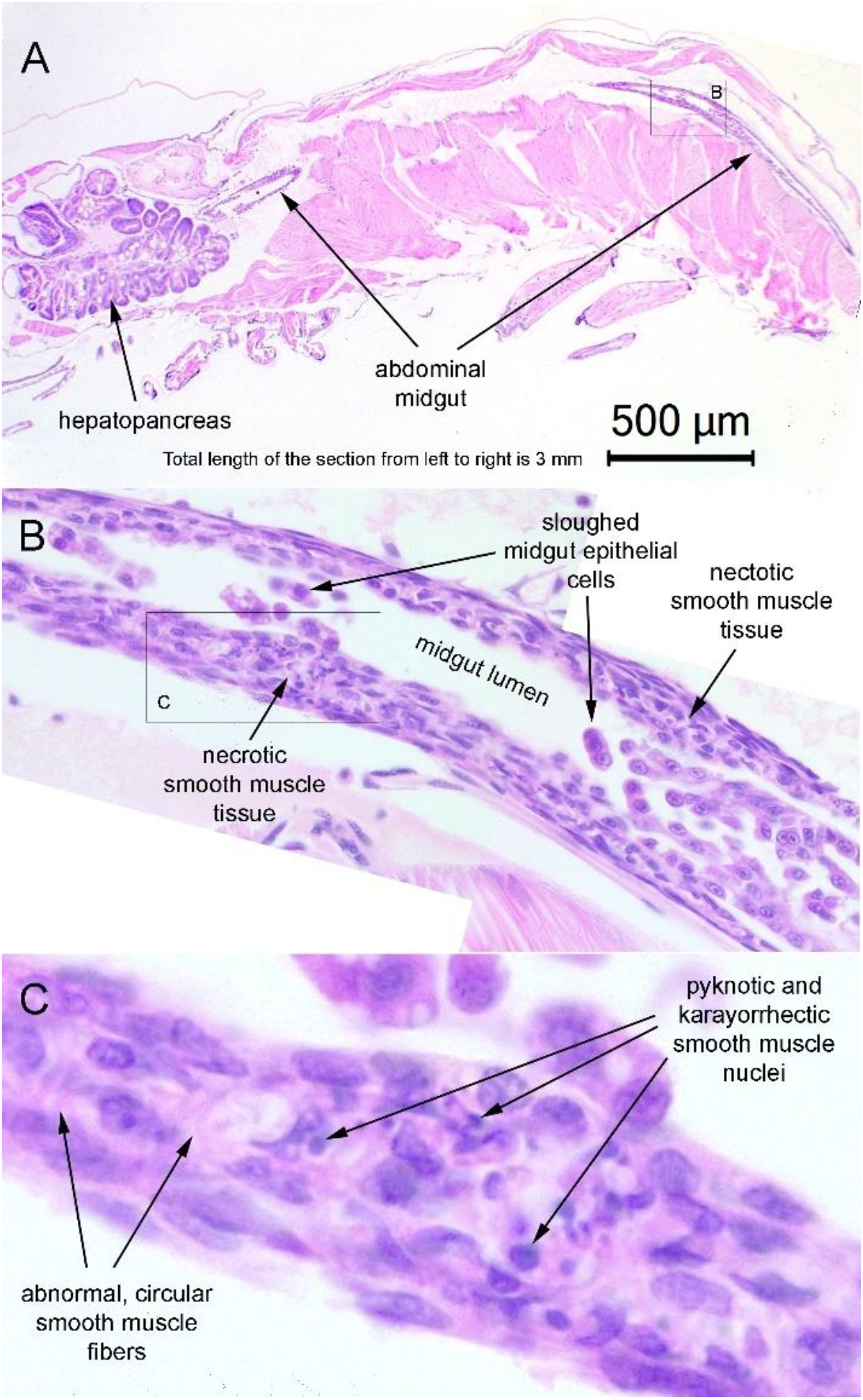
Photomicrographs of a PL specimen from Tank A showing MSMN in a lateral section of the midgut along the longitudinal axis. The 6^th^ abdominal segment is not in the cut. Diagnosis requires high magnification and does not give as clear an indication of the severity of muscle necrosis as does the tangential section in previous Fig. 5.

Moreover**, Fig. 7** shows MSMN histopathology of a third PL specimen from hatchery Tank A. Again, the 6^th^ abdominal segment is not in the cut. The whole tissue section (**Fig. 7A**) includes both cross and tangential sections of the midgut in its mid-abdominal region and a cross-section of the midgut region near its exit from within the HP. MSMN is apparent in the mid-abdominal region of the midgut (**Fig. 7B, C**), but not in the midgut region near the hepatopancreas (**Fig. 7D, E**). The reason for the specificity of MSMN pathology to the midabdominal region of the PL body suggests that the pathology may originate in the mid-abdominal region and then spread from there in both anterior and posterior directions. In any case, our detection of MSMN was more successful in this region, suggesting that one should focus on examining the midgut in its mid-abdominal region for MSMN diagnosis.

**Figure 7.**
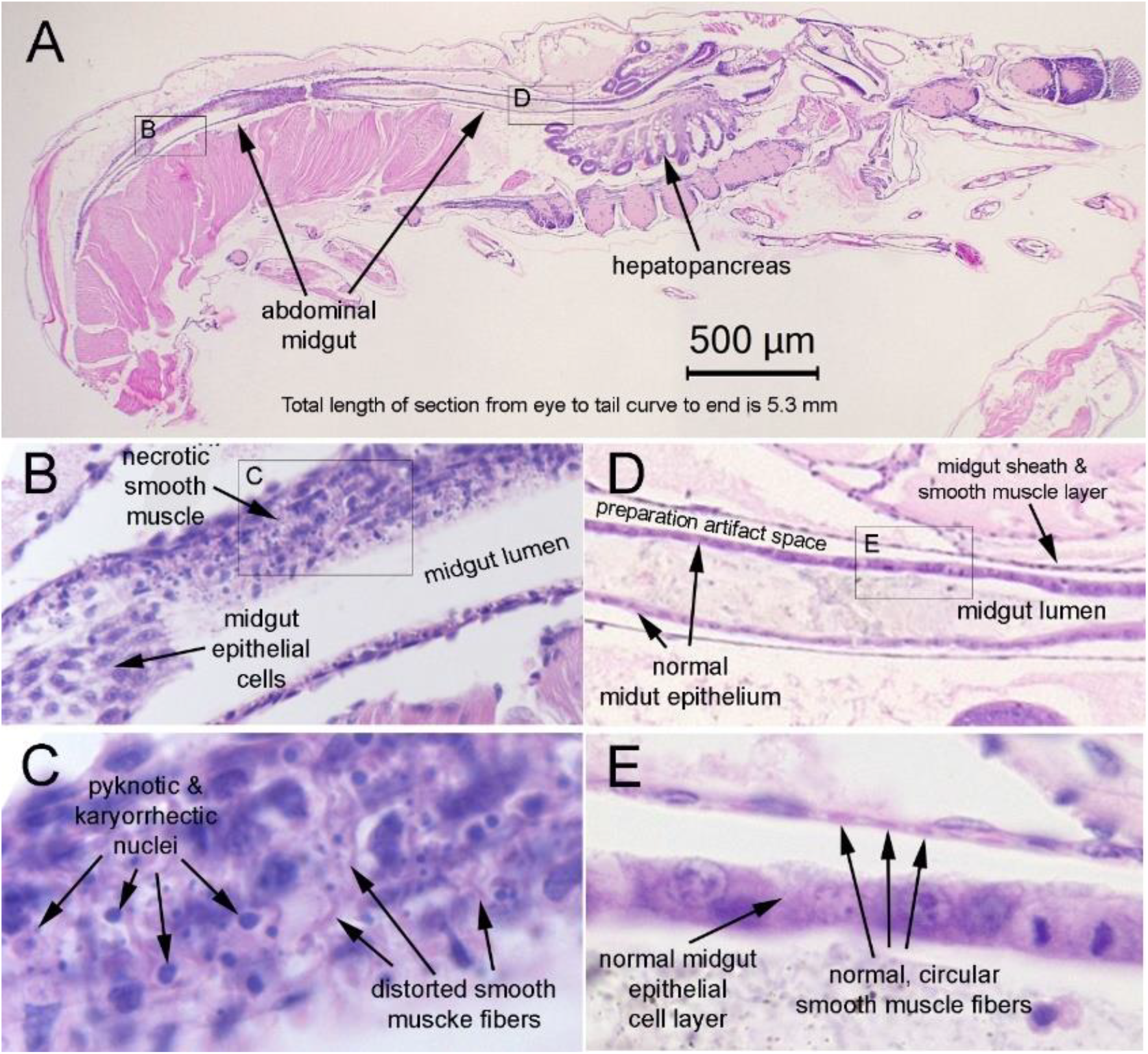
Photomicrographs of a PL specimen from hatchery Tank A showing MSMN necrosis in the mid-abdominal portion of the midgut, but normal histology in the portion of the midgut near the hepatopancreas. (A) Whole specimen slice with the 6^th^ abdominal segment missing from the cut. (B) Mid-abdominal midgut section showing epithelial cells together with overlying necrotic smooth muscle tissue (MSMN). (C) High magnification of the severe muscle necrosis characterized by pyknotic and karyorrhectic nuclei plus distorted, eosinophilic, smooth muscle fibers resembling cooked spaghetti noodles. (D) Section of the midgut near the HP showing normal histology that contrasts with the necrosis in the mid-abdominal region.

An additional specimen from hatchery Tank A, representing a fourth PL individual, exhibited MSMN histopathology as shown in **Fig. 8**. Again the 6^th^ abdominal segment is not within the cut. High magnification images in Fig. 8C and D were obtained by changing the location and level of focus to obtain sharp images on the normal midgut epithelial cells (8C) and the overlying necrotic smooth muscle tissue (8D). The latter clearly shows pyknotic and karyorrhectic nuclei and distorted smooth muscle fibers characteristic of MSMN.

**Figure 8.**
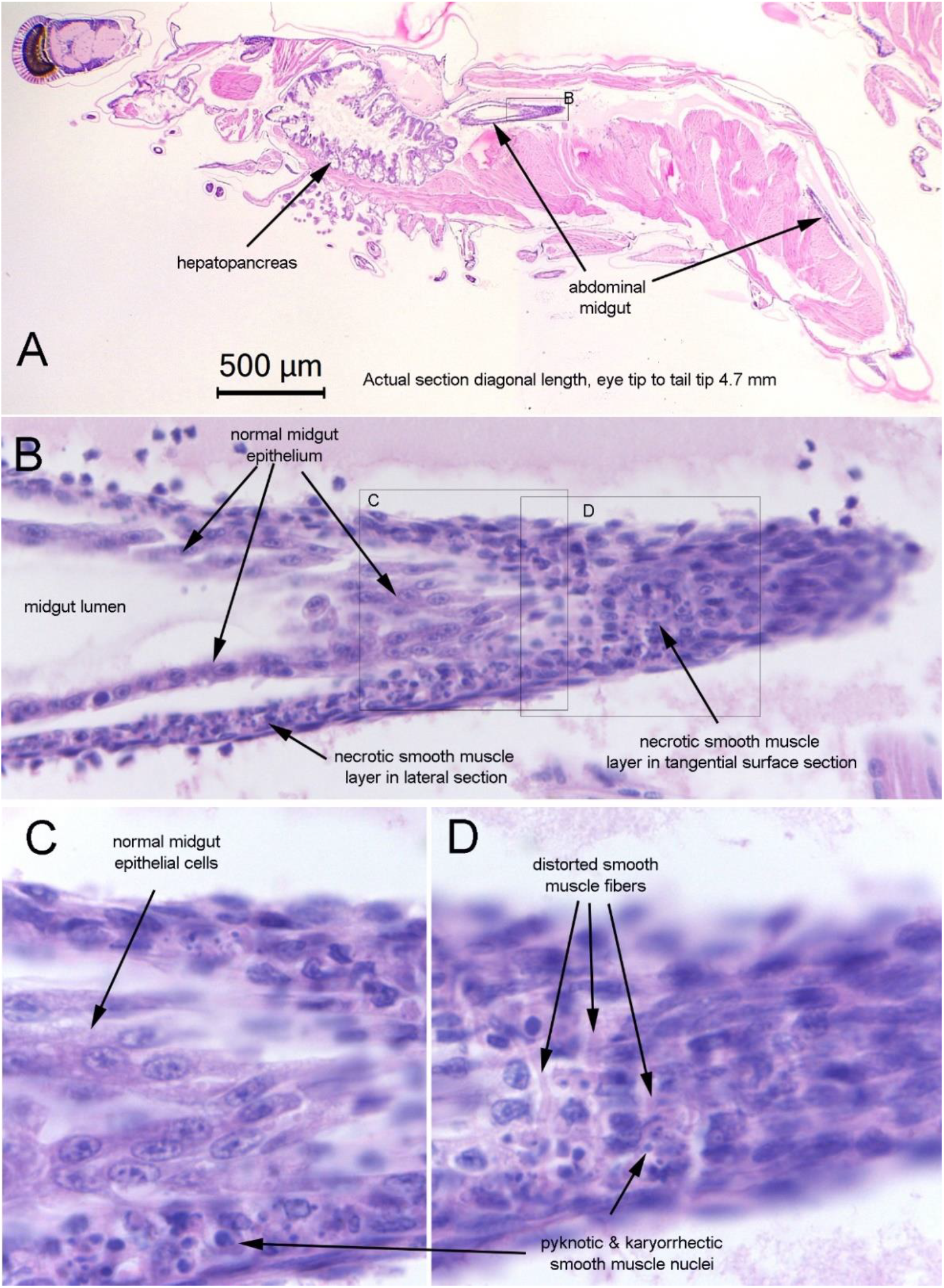
Photomicrographs of a 3rd PL specimen from hatchery Tank A with a tangential cut through a portion of the midgut in the 1^st^ abdominal segment. This includes both lateral sections and tangential sections through the midgut again revealing the need for tangential cuts to clearly see the histopathology of MSMN.

From hatchery Tank B, the section of only one specimen showed MSMN pathology, as shown **Fig. 9**. However, the cut did not include the hepatopancreas. In addition, the cut through the midgut wall was mostly longitudinal, showing the smooth muscle layers in cross section, similar to the cut shown in Fig. 7 for the specimen from Tank A.

**Figure 9.**
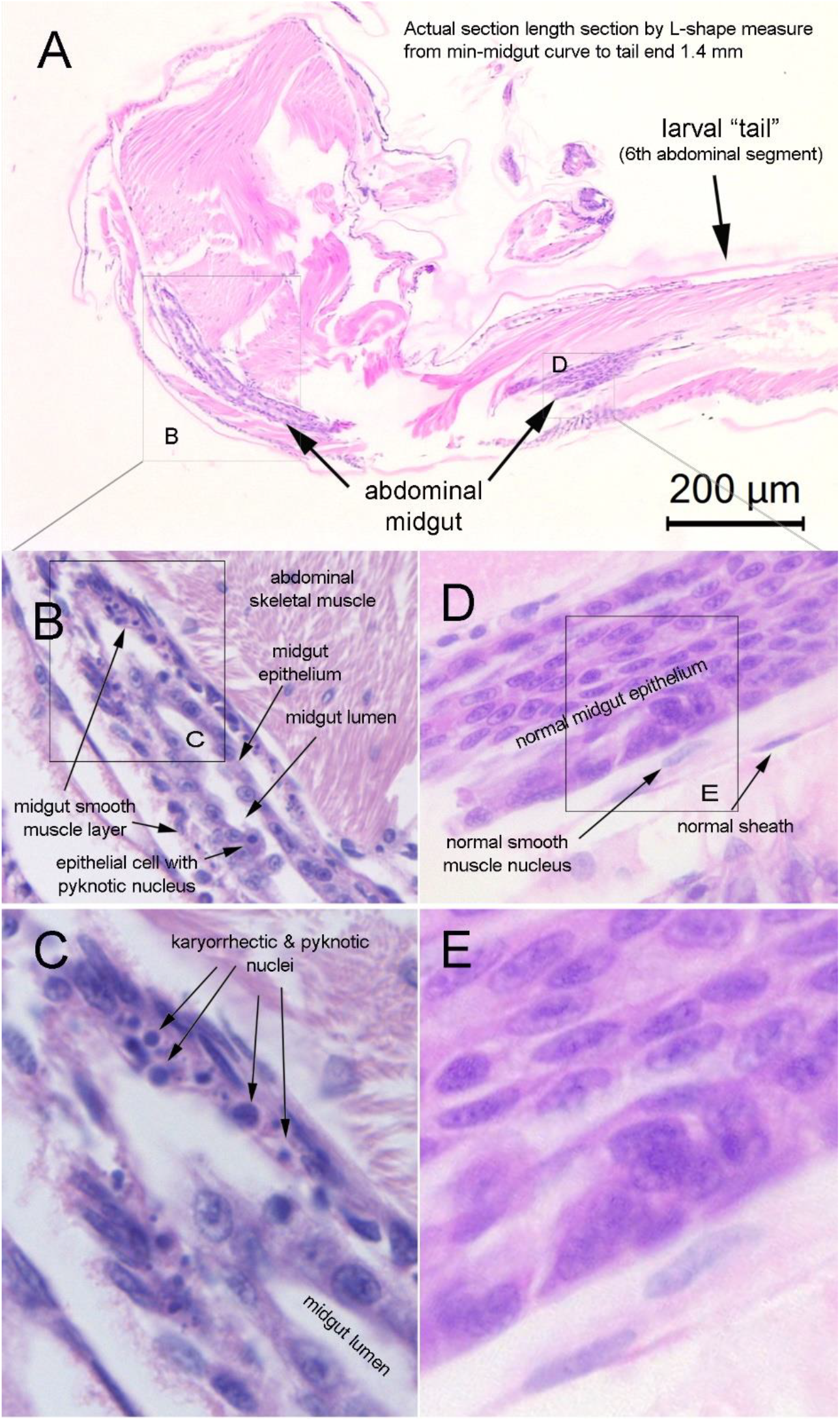
Photomicrographs of the single specimen from Tank B that exhibited MSMN pathology. (A) Whole larval section with cephalothorax missing but exhibiting MSMN in the midgut in the mid-abdominal region but with normal midgut in the 6^th^ abdominal segment. (B, C) Magnified region from box B in A, rotated slightly to increase the length of the midgut exhibiting MSMN pathology. (D, E) Magnified region from box D in A.

To summarize the histopathology of MSMN, it appears to originate in smooth muscles in the wall of the PL midgut where it is located within the first 5 segments of the PL abdomen. This excludes the midgut region within the cephalothorax near the hepatopancreas and the midgut in the 6^th^ abdominal segment that is disproportionally extended in PL into a “tail” that contains the hindgut at its end near the telson. The PL midgut itself is in the range of about 1-2 mm in diameter with a wall less than 10 microns in thickness that contains 4 cell layers.

These include the midgut epithelium surrounding the lumen followed by an underlying circular smooth muscle layer, then a longitudinal smooth muscle layer and finally a sheath. Thus, each layer is only 1-3 microns thick, so that the frequency of tangential tissue slices such as those shown here in Figs.5,7and 8 that give a surface view of the smooth muscle layer would be rare.

Due to these constraints, we believe that MSMN lesions may have been previously overlooked because the abdominal midgut is not included in the cephalothorax tissue sections often used for routine shrimp histopathological analysis. In addition, when the abdomen is included, the tissue sections are mid-lateral cuts (as with the whole PL herein) in which the midgut appears as two relatively thin lines of tissue on either side of the gut lumen. Each side consists of midgut epithelial cells that sit on a relatively thin layer of underlying muscle tissue that, in turn, is bound by connective tissue that forms the gut sheath. This can be seen here best in Figs. 4 to 7 and compared to the portion of the midgut that has been cut tangentially to its long axis. The latter allows a much better and more detailed planal view of the MSMN pathology in the muscle tissue layer (less commonly seen in such sections). Even so, the pathology is very difficult to detect using a microscope unless at least a 40x objective is used and special attention is paid to the midgut in the mid abdominal region.

## DISCUSSION

Our histological analysis has revealed MSMN lesions in two hatchery tanks exhibiting TPD/GPD, suggesting that it may be a histopathological anomaly associated with the disease. We found no bacteria or parasites in these lesions. Thus, it is possible that MSMN lesions could arise in response to some type of environmental toxin(s) or microbial toxin(s) such as the non-Pir-like toxins produced by the stains of *Vibrio parahaemolyticus* recently proposed to be the cause of TPD (Liu *et al.*, 2023; Zou *et al.*, 2020). The positive PCR detection of *vhvp*-2 toxin were found in the specimens from both tanks suggesting that the *vhvp-2* toxin might be involved with MSMN lesions. The effect of purified *vhvp-2* toxin on smooth muscle necrosis should be confirmed and the information might lead to a practical application to control MSMN-associated translucence in PL. Alternatively, they may be caused by the newly proposed *Baishivirus* (Xu *et al.*, 2023). Either way, proof of a causal link between TPD/GPD and MSMN would provide a method for detection of TPD/GPD using standard histological methods. Any archived samples of TPD/GPD specimens in paraffin blocks or available as H&E-stained slides can be retrospectively examined for the presence of MSMN lesions. Currently, the lack of published information regarding the histopatholgy of PL exhibiting GPD, and especially the lack of *in situ* hybridization assays to determine the viral target tissue make it impossible to distinguish between TPD and GPD at the gross level.

With respect to field detection of MSMN lesions using fresh material, it may be possible to prepare small portions of mid-abdominal midgut tissue specifically removed for rapid fixation and H&E staining in 2.5 ml Eppendorf tubes followed by preparation of squash mounts in the same manner as whole gill mount fragments have been used for detection of yellow head virus (YHV) and white spot syndrome virus (WSSV) (Flegel *et al.*, 1997). This might aid in more rapid diagnosis of MSMN in the field where PCR technology may not be available. At the same time, since MSMN must prevent paristalsis, it may be possible by using a sufficiently powerful handlens to examine living PL from a hatchery tank for absence of peristalsis as an early warning for transparent PL. In addition, preparation of fresh specimens fixed in a manner to preserve RNA would allow for *in situ* hybridization tests for the presence of *Baishivirus* in MSMN lesions, should the virus be present. Ideally, laboratory challenges using purified *Baishivirus* with appropriate controls would settle this question. We are willing to cooperate with colleagues anywhere in the Asia-Pacific region to explore these possibilities.

## Acknowledgement

This research was supported by the National Science, Research and Innovation Fund, Thailand Science Research and innovation (TSRI) (Grant No: FFB680075/0337) and received funding support from the NSRF via the Program Management Unit for Human Resources & Institutional Development, Research and Innovation (Grant No: B05F640137).

